# Combinatorial morphogenetic and mechanical cues to mimic bone development for defect repair

**DOI:** 10.1101/561837

**Authors:** S. Herberg, A. M. McDermott, P. N. Dang, D. S. Alt, R. Tang, J. H. Dawahare, D. Varghai, J-Y. Shin, A. McMillan, A. D. Dikina, F. He, Y. Lee, Y. Cheng, K. Umemori, P.C. Wong, H. Park, J. D. Boerckel, E. Alsberg

## Abstract

Endochondral ossification during long bone development and natural fracture healing initiates by mesenchymal cell condensation and is directed by local morphogen signals and mechanical cues. Here, we aimed to mimic these developmental conditions for regeneration of large bone defects. We hypothesized that engineered human mesenchymal stem cell (hMSC) condensations with *in situ* presentation of transforming growth factor-β1 (TGF-β1) and/or bone morphogenetic protein-2 (BMP-2) from encapsulated microparticles would promote endochondral regeneration of critical-sized rat femoral bone defects in a manner dependent on the *in vivo* mechanical environment. Mesenchymal condensations induced bone formation dependent on morphogen presentation, with dual BMP-2 + TGF-β1 fully restoring mechanical bone function by week 12. *In vivo* ambulatory mechanical loading, initiated at week 4 by delayed unlocking of compliant fixation plates, significantly enhanced the bone formation rate in the four weeks after load initiation in the dual morphogen group. *In vitro*, local presentation of either BMP-2 alone or BMP-2 + TGF-β1 initiated endochondral lineage commitment of mesenchymal condensations, inducing both chondrogenic and osteogenic gene expression through SMAD3 and SMAD5 signaling. *In vivo*, however, endochondral cartilage formation was evident only in the BMP-2 + TGF-β1 group and was enhanced by mechanical loading. The degree of bone formation was comparable to BMP-2 soaked on collagen but without the ectopic bone formation that limits the clinical efficacy of BMP-2/collagen. In contrast, mechanical loading had no effect on autograft-mediated repair. Together, this study demonstrates a biomimetic template for recapitulating developmental morphogenic and mechanical cues *in vivo* for tissue engineering.

**One Sentence Summary:** Mimicking aspects of the cellular, biochemical, and mechanical environment during early limb development, chondrogenically-primed human mesenchymal stem cell condensations promoted functional healing of critical-sized femoral defects via endochondral ossification, and healing rate and extent was a function of the *in vivo* mechanical environment.

## Introduction

Endochondral ossification is an indirect mode of bone formation that occurs during long bone development and natural fracture repair whereby mesenchymal progenitor cells form a cartilage anlage that is replaced by bone (*1, 2*). In both development and repair, mechanical cues are essential for proper endochondral ossification. For example, experimental fetal paralysis significantly decreases bone mass *in ovo* (*3*) and motion *in utero* is important for normal bone and joint development (*4*). Likewise, during fracture repair, the amount and mode of interfragmentary strain determines whether a fracture will heal through endochondral or intramembranous ossification (*3, 5-11*). Though bone fractures heal with 90-95% success rates by forming a cartilaginous callus (*12-14*), large bone defects greater than 3 cm in length cannot form a callus and exhibit high complication rates, representing a significant clinical burden (*15*). Current standard treatments for large bone defects include autologous bone grafting and delivery of high-dose recombinant human bone morphogenetic protein-2 (BMP-2) soaked on a collagen sponge carrier; however, these treatments are limited by donor-site morbidity and ectopic bone formation, respectively (*16, 17*).

Cell-based tissue engineering strategies may provide a promising alternative to bone grafts. One proposed strategy combines osteogenic/progenitor cells with materials that mimic the structural properties of mature bone. However, poor cell engraftment and viability due to insufficient vascular supply limit the efficacy of osteogenic cell delivery (*18-21*), and the rigidity of mature bone matrix-like scaffolds can impede stimulatory mechanical loads (*22*). An alternative strategy is to seek to mimic the process by which bone tissue forms during development, namely endochondral ossification (*23-29*). As the cartilage anlage is mechanically compliant, avascular, and capable of naturally recruiting neovasculature and endogenous progenitors and osteoblasts, this approach may overcome key challenges for the regeneration of challenging bone defects. Here, we sought to recapitulate the (1) mesenchymal condensation, (2) sequential morphogenetic cues, and (3) mechanical cues that mediate developmental endochondral ossification for regeneration of critical-sized bone defects in adult rats.

Mesenchymal cell condensation and chondrogenic differentiation in the developing limb bud are regulated by sequential morphogenic signals, including TGF-β (*30*) and BMP (*31*), which mediate cell condensation and induce the master chondrogenic transcription factors SRY-Box (SOX) 5, 6, and 9 (*32, 33*). Recent studies have shown that avascular cartilage templates derived from hMSC aggregates are capable of progressing through endochondral ossification (*25, 26, 29, 34-37*), producing mineralized matrix, vasculature, and a bone marrow hematopoietic stem cell niche (*38*), but these required extended pre-culture with exogenous growth factors *in vitro* for chondrogenic priming. To address this problem, we developed scaffold-free hMSC condensations, formed through self-assembly of cellular sheets incorporated with TGF-β1-releasing gelatin microspheres for *in situ* chondrogenic priming. These formed robust cartilage tissue *in vitro* (*39*) and induced endochondral bone defect regeneration after implantation *in vivo* (*40*). Further, sustained individual or co-delivery of BMP-2 in hMSC condensations induced both chondrogenesis and osteogenesis *in vitro* (*41, 42*) and endochondral regeneration of calvarial defects *in vivo* (*27*). Local morphogen presentation circumvents the need for lengthy exogenous supplementation of inductive signals and enables *in vivo* implantation of the cellular constructs in a timely and cost-efficient manner, thereby providing a promising system to investigate the progression of mesenchymal condensation through endochondral ossification *in vivo.*

The mechanical environment considerably influences bone development, homeostasis, and regeneration (*3, 5-11*). To investigate the roles of mechanical cues in large bone defect regeneration, we developed a critically-sized rat femoral bone defect model in which ambulatory load transfer can be controlled by dynamic modulation of axial fixation plate stiffness (*40, 43-45*). We previously showed that load initiation, delayed to week four (after the onset of regeneration and bony bridging), significantly enhanced bone formation, biomechanical properties, and local tissue adaptation mediated by BMP-2-releasing hydrogels (*43-45*). Recently, we showed that *in vivo* loading of engineered hMSC condensations, containing TGF-β1-releasing gelatin microparticles, restored bone function through endochondral ossification (*40*).

However, these studies focused on single morphogen presentation, and bone development features an intricate and tightly coordinated sequence of both morphogenetic and mechanical cues. Here, we investigated the combinatorial roles of controlled temporal presentation of TGF-β1 and/or BMP-2, to mimic events in the developing limb bud, with *in vivo* mechanical loading. To control local morphogen presentation, we engineered mesenchymal condensations incorporated with gelatin or mineral microparticles for local release of TGF-β1 to drive chondrogenesis and BMP-2 to promote remodeling of the cartilaginous anlage to bone, respectively. To regulate *in vivo* mechanical conditions, we dynamically modulated fixation plate stiffness, altering ambulatory load-sharing. We found that morphogen co-delivery and *in vivo* mechanical loading combinatorially regulated bone regeneration and directed ossification mode, with combined treatment inducing full functional restoration of bone mechanical properties.

## Results

### Effects of in vivo mechanical loading on autograft-mediated bone regeneration

Previously, we (*40, 44, 45*) and others (*46, 47*), found that *in vivo* mechanical loading can enhance the regeneration of large bone defects. First, to test whether loading can enhance autograft-mediated bone regeneration, we treated critical-sized (8 mm) defects in Rowett nude (RNU) rat femora with morselized cortical bone autografts in two groups: control (stiff fixation plates) and delayed loading (compliant plates, unlocked to allow ambulatory load transfer at week 4) (Supplementary Figure S1A). Bone formation was evaluated over 12 weeks by longitudinal microcomputed tomography (microCT) imaging. Mechanical loading did not affect autograft-mediated bone formation (Supplementary Figure S1B,C). Since stiff load-bearing scaffolds can impede load-induced bone repair (*22*), we next tested whether non-load bearing mesenchymal condensations containing local presentation of BMP-2 could promote bone defect repair.

### Comparison of BMP-2-containing mesenchymal condensations with the current standard of care in absence of mechanical loading

To this end, we compared the bone formation capacity of BMP-2-containing mesenchymal condensations with either autograft or BMP-2-loaded collagen sponge controls, without mechanical loading (i.e., stiff fixation). hMSC condensations were assembled with mineral-coated hydroxyapatite microparticles for *in situ* controlled presentation of 2 μg BMP-2 (*27, 41, 42, 48, 49*). The BMP-2/collagen group received the same dose of BMP-2 (2 μg), adsorbed onto lyophilized collagen sponges, and the autograft group featured morselized cortical bone.

High-resolution *ex vivo* microCT analysis was performed at week 12 to evaluate bone formation and architecture. Both BMP-2/collagen and morselized autograft produced significantly greater bone volume fraction and trabecular number, and smaller trabecular separation compared to BMP-2-containing hMSC condensations (Supplementary Figure S2A-E). However, ectopic bone formation (i.e., bone extending beyond the 5-mm defect diameter) was significantly greater in defects treated with BMP-2 delivered on collagen compared to BMP-2-containing hMSC condensations (∼3-fold) or autografts (∼4-fold; Supplementary Figure S2A,F). While the development-mimetic mesenchymal condensations induced bone formation with less ectopic bone than BMP-2/collage, their ultimate regenerative capacity was inferior.

### Controlled combinatorial morphogen presentation with in vivo mechanical load transfer

However, natural bone development and fracture repair occur through endochondral ossification in response to combined chondrogenic, osteogenic, and mechanical cues. Therefore, we hypothesized that these factors would be required in a combinatorial fashion to induce bone regeneration in manner that reproduces natural bone formation. To this end, we next treated defects with hMSC condensations containing local presentation of TGF-β1 and/or BMP-2, with or without *in vivo* mechanical loading.

We previously showed that delayed *in vivo* mechanical loading, initiated at week 4 by compliant fixation plate unlocking, moderately enhanced (18%) bone regeneration by cell-free hydrogel-delivered BMP-2 (*44, 45*) and substantially enhanced (180%) endochondral regeneration by TGF-β1-incorporated mesenchymal condensations (*40*). Here, we investigated the interactions of mechanical loading with morphogen presentation. TGF-β1 was presented in gelatin microparticles for early release and BMP-2 from hydroxyapatite microparticles for sustained release (*39*) Three morphogen conditions were evaluated: human mesenchymal condensations with empty microparticles (empty controls), condensations with BMP-2-releasing microparticles, and condensations with BMP-2 + TGF-β1-releaseing microparticles. Each was implanted *in vivo* with either stiff plates as (control) or compliant plates unlocked at week 4 (delayed loading), for a total of six groups (Table 1).

**Table 1.** Experimental Design.

| **Group** | **Morphogen Condition** | **Mechanical Loading Condition** |
| --- | --- | --- |
| 1 | Empty/Control | Stiff |
| 2 | BMP-2 | Stiff |
| 3 | BMP-2 + TGF-β1 | Stiff |
| 4 | Empty/Control | Delayed (compliant plate unlocked at week 4) |
| 5 | BMP-2 | Delayed (compliant plate unlocked at week 4) |
| 6 | BMP-2 + TGF-β1 | Delayed (compliant plate unlocked at week 4) |

### In vivo radiography and microCT analyses

Longitudinal X-ray radiography and microCT analyses were performed at weeks 4, 8, and 12. BMP-2-containing hMSC condensations enhanced bone regeneration relative to controls at weeks 8 and 12 with both loading regimens (Figure 1A,B red lines/boxes, C). Some instances of bridging were observed (stiff: 3/9; compliant: 4/7; Supplementary Figure S3). The BMP-2-mediated regenerative effects were significantly enhanced by TGF-β1 co-delivery (Figure 1A,B blue lines/boxes, C). Bridging was achieved in nearly all dual-morphogen samples (stiff: 9/11; compliant: 9/10; Supplementary Figure S3).

**Figure 1:**
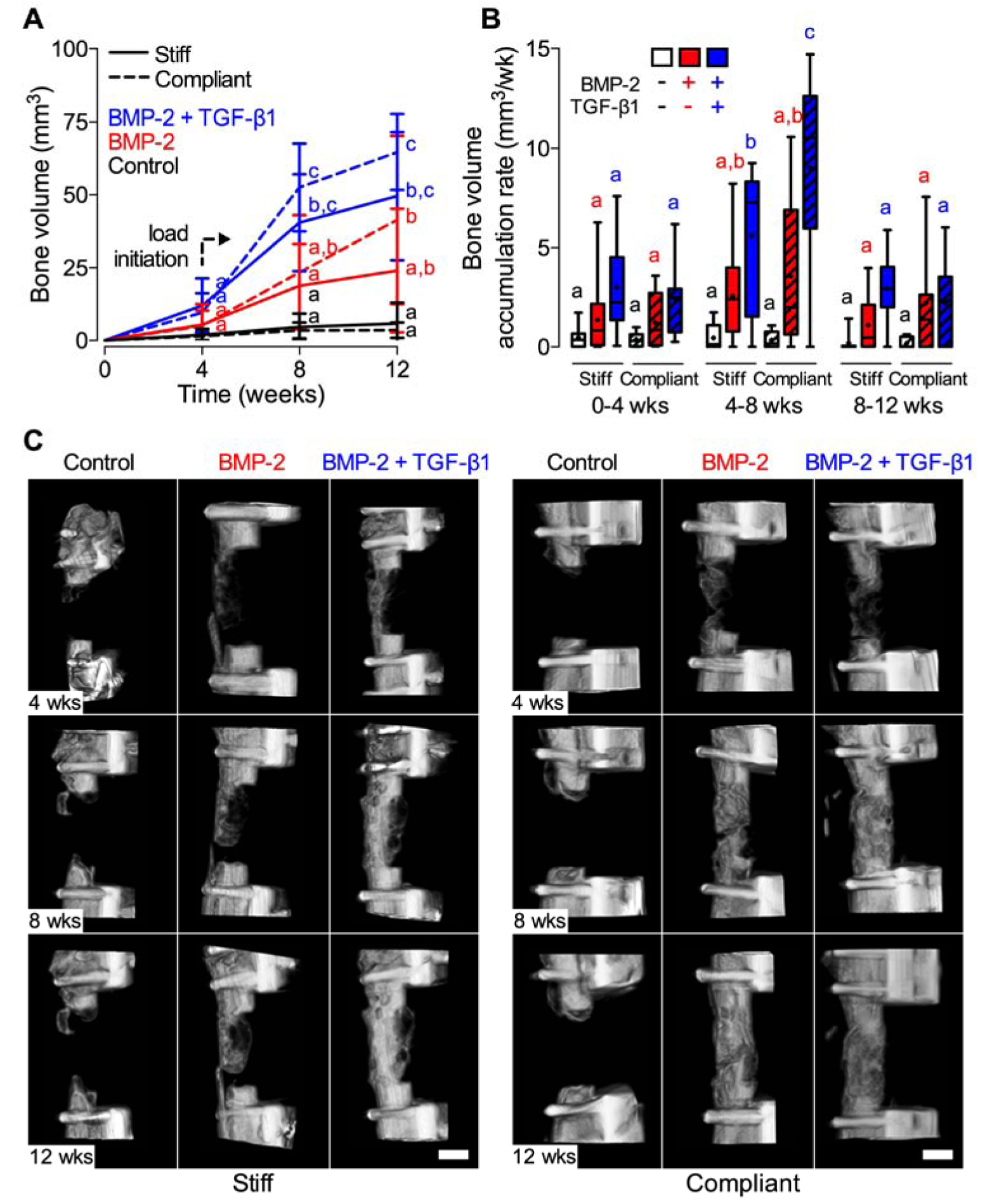
Effects of morphogen priming of engineered hMSC condensations and *in vivo* mechanical loading on longitudinal bone formation and bone accumulation rate. (**A**) Longitudinal quantification of bone volume at weeks 4, 8, and 12 by *in vivo* microCT (N = 4-11 per group). Data shown with mean ± SD. (**B**) Bone volume accumulation rate, defined as bone volume accrual over each 4-week interval. Box plots display median as horizontal line, mean as +, inter-quartile range as boxes, and min/max range as whiskers. (**C**) Representative 3-D microCT reconstructions showing bone formation per group over time. Representative samples were selected based on mean bone volume at week 12. Scale bar, 3 mm. Comparisons between groups were evaluated by two-way repeated measures ANOVA with Tukey’s post-hoc tests. Repeated significance indicator letters (a,b,c) signify p > 0.05, while groups with distinct indicators signify p < 0.05 at each time point. Comparisons between time points were not assessed.

Mechanical loading significantly elevated the bone volume accumulation rate during the four weeks immediately after compliant plate unlocking in BMP-2 + TGF-β1-presenting condensations compared to all other groups and time intervals (Figure 1B). New bone formation was negligible in empty control samples, regardless of mechanical loading (Figure 1A,B black lines/boxes, C), and none achieved bridging by week 12 (stiff: 0/8; compliant: 0/4; Supplementary Figure S3).

Thus, transplanted hMSC condensations induced bone regeneration dependent on morphogen identity, and mechanical loading influenced the rate of bone formation during the four weeks following load initiation in samples containing both BMP-2 and TGF-β1.

### Ex vivo microCT analysis

Tissue composition and organization was then evaluated at high-resolution by *ex vivo* microCT analysis at week 12. Control hMSC condensations without morphogen presentation failed to induce healing regardless of mechanical loading, with new bone formation merely capping the exposed medullary canals, predominantly on the proximal end (Figure 2; Supplementary Figure S4). Both BMP-2 and BMP-2 + TGF-β1 presentation enhanced bone regeneration compared to control condensations (Figure 2A,B). New bone within the defects exhibited an approximately uniform proximal-to-distal distribution (Supplementary Figure S4A), and lacked notable ectopic bone (Supplementary Figure S4B), in contrast to BMP-2 delivery on collagen sponge (Supplementary Figure 1). Dual morphogen presentation and mechanical loading together produced regenerated bone with a trabecular internal architecture contained within a cortical shell, quantitatively similar in structure to native trabecular/cortical bone architecture (Figure 2C-E; Supplementary Figure S4C-F). These data show that bone distribution and architecture were determined primarily by presented morphogen identity.

**Figure 2:**
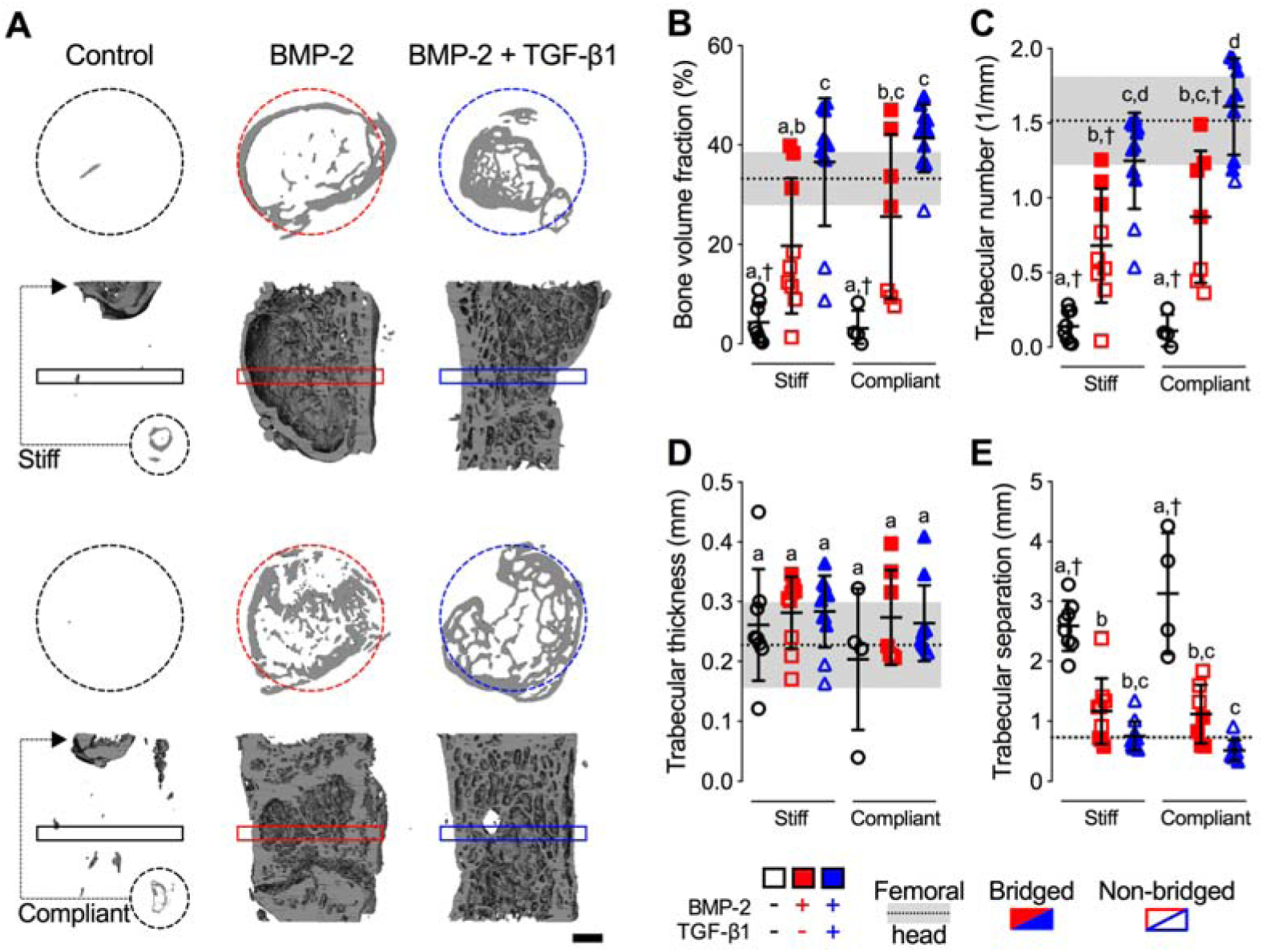
Effects of morphogen priming of engineered hMSC condensations and *in vivo* mechanical loading on new bone quantity and architecture. (**A**) Representative 3-D microCT reconstructions, with bone formation illustrated at mid-shaft transverse (top) and sagittal (bottom) sections at week 12, selected based on mean bone volume. Dashed circles show 5 mm defect region. Rectangular boxes illustrate transverse cutting planes. Note, due to minimal bone regeneration, additional transverse sections for stiff and compliant no growth factor controls were derived from the proximal end of the defect (small dashed circles and arrows). Scale bar, 1 mm. (**B**) Morphometric analysis of bone volume fraction, (**C**) trabecular number, (**D**) trabecular thickness, and (**E**) trabecular separation at week 12 (N = 4-11 per group), shown with corresponding measured parameters of femoral head trabecular bone (N = 3; dotted lines with gray shading: mean ± SD; ^†^ p<0.05 vs. femoral head). Individual data points shown with mean ± SD. Comparisons between groups were evaluated by two-way ANOVA with Tukey’s post-hoc tests. Repeated significance indicator letters (a,b,c) signify p > 0.05, while groups with distinct indicators signify p < 0.05.

### Restoration of mechanical bone function

Next, we evaluated the restoration of limb mechanical properties by torsion testing to failure at week 12, in comparison to age-matched intact femurs. BMP-2 + TGF-β1-containing hMSC condensations enhanced stiffness and failure torque compared to controls. Mechanical loading did not significantly alter mechanical properties compared to corresponding stiff plate controls for each group, but significantly increased the mean polar moment of inertia (a measure of structural cross-sectional geometry) and fully restored functional mechanical properties in the BMP-2 + TGF-β1 group (Figure 3A-C), with statistically equivalent torsional stiffness and maximum torque at failure compared to age-matched intact femurs (Figure 3A,B cf. gray band).

**Figure 3:**
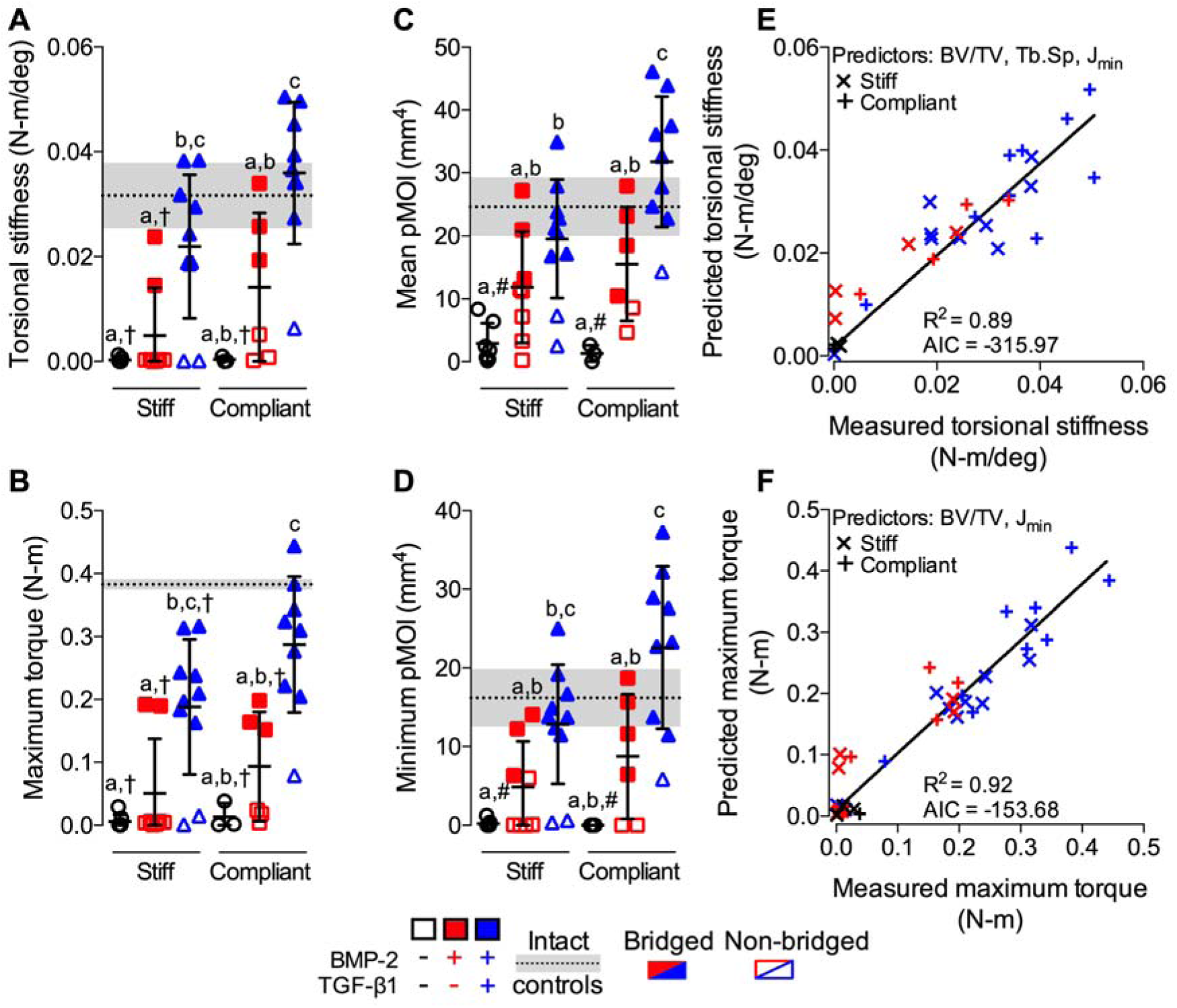
Effects of morphogen priming of engineered hMSC condensations and *in vivo* mechanical loading on functional defect regeneration. (**A**) Torsional stiffness, (**B**) maximum torque at failure, (**C**) mean polar moment of inertia (pMOI), and (**D**) minimum pMOI. Best subsets regression analysis (R^2^) with lowest Akaike’s information criterion (AIC) value for measured and predicted (**E**) torsional stiffness and (**F**) maximum torque at failure indicating significant contributions of bone volume fraction (BV/TV), trabecular separation (Tb.Sp), and minimum pMOI (J_min_). Individual data points shown with mean ± SD (N = 3-10 per group). Comparisons between groups were evaluated by two-way ANOVA with Tukey’s post-hoc tests. Repeated significance indicator letters (a,b,c) signify p > 0.05, while groups with distinct indicators signify p < 0.05. Biomechanical and structural parameters are shown with age-matched intact bone properties, with pMOI obtained from the same midshaft ROI as used for the defects (N = 3; dotted lines with gray shading: mean ± SD; ^†,#^p<0.05 vs. intact bone).

To identify the key structural predictors of mechanical behavior (*50*), we performed a type II multivariate best-subsets regression analysis with model predictors selected by minimization of the Akaike’s information criterion (AIC) (*51*). For torsional stiffness, bone volume fraction, trabecular separation, and minimum pMOI were the best combined predictors (Figure 3E; Supplementary Figure S5A-D). For maximum torque, bone volume fraction and minimum pMOI were the best combined predictors (Figure 3F; Supplementary Figure S5E-H). Thus, the mechanical properties were determined by the amount, distribution, and trabecular organization of the regenerate bone.

Together, these data indicate that restoration of biomechanical competence was dependent on the identity of presented morphogens and induced full functional repair only by dual morphogen presentation with *in vivo* mechanical loading.

### In vitro signaling and differentiation analyses

We hypothesized that the cellular organization into condensations and the development-mimetic morphogen presentation would induce endochondral bone formation. TGF-β superfamily ligands bind to type I and II serine/threonine kinase receptor complexes and transduce signals via SMAD proteins (*52*). In the developing limb, TGF-β signaling has been shown to occur early during the chondrogenic cascade, prior to the BMPs (*30, 32, 33*). Further, a recent study proposed that transient activation of the TGF-β pathway may be required to promote a chondrogenic response to BMP signaling during later stages of chondrogenesis (*53*). Therefore, we next tested the effects of combinatorial morphogen presentation on chondrogenic and osteogenic activity of the mesenchymal condensations in vitro. As prepared for *in vivo* transplantation, TGF-β1 was presented in an early manner by release from gelatin microspheres, while BMP-2 was released in a more sustained manner from mineral-coated hydroxyapatite microparticles.

After two days *in vitro* culture (coinciding with the time of *in vivo* transplantation), the engineered hMSC condensations exhibited homogeneous cellular organization across groups, with uniformly distributed microspheres and no detectable GAG or mineral deposition (Figure 4A, Supplementary Figure S6A). Transcript analysis of key differentiation markers revealed that either TGF-β1 or BMP-2 presentation alone significantly induced mRNA expression of genes indicative of both chondrogenic (SOX9, aggrecan (ACAN), and collagen type 2A1 (COL2A1)) and osteogenic (ALP) priming, relative to growth factor-free controls (Figure 4B, Supplementary Figure S6B). BMP-2 + TGF-β1 co-delivery further increased expression of SOX9, ACAN, COL2A1, ALP and osterix (OSX) mRNA (Figure 4B). Lastly, BMP-2 presentation significantly increased both SMAD3 and SMAD5 phosphorylation relative to control condensations without growth factor, and was significantly potentiated by TGF-β1 co-delivery (Figure 4C,D). These *in vitro* data suggest that presentation of either BMP-2 or BMP-2 + TGF-β1 induced chondrogenic lineage priming via both SMAD3 and SMAD5 signaling at the time on implantation.

**Figure 4:**
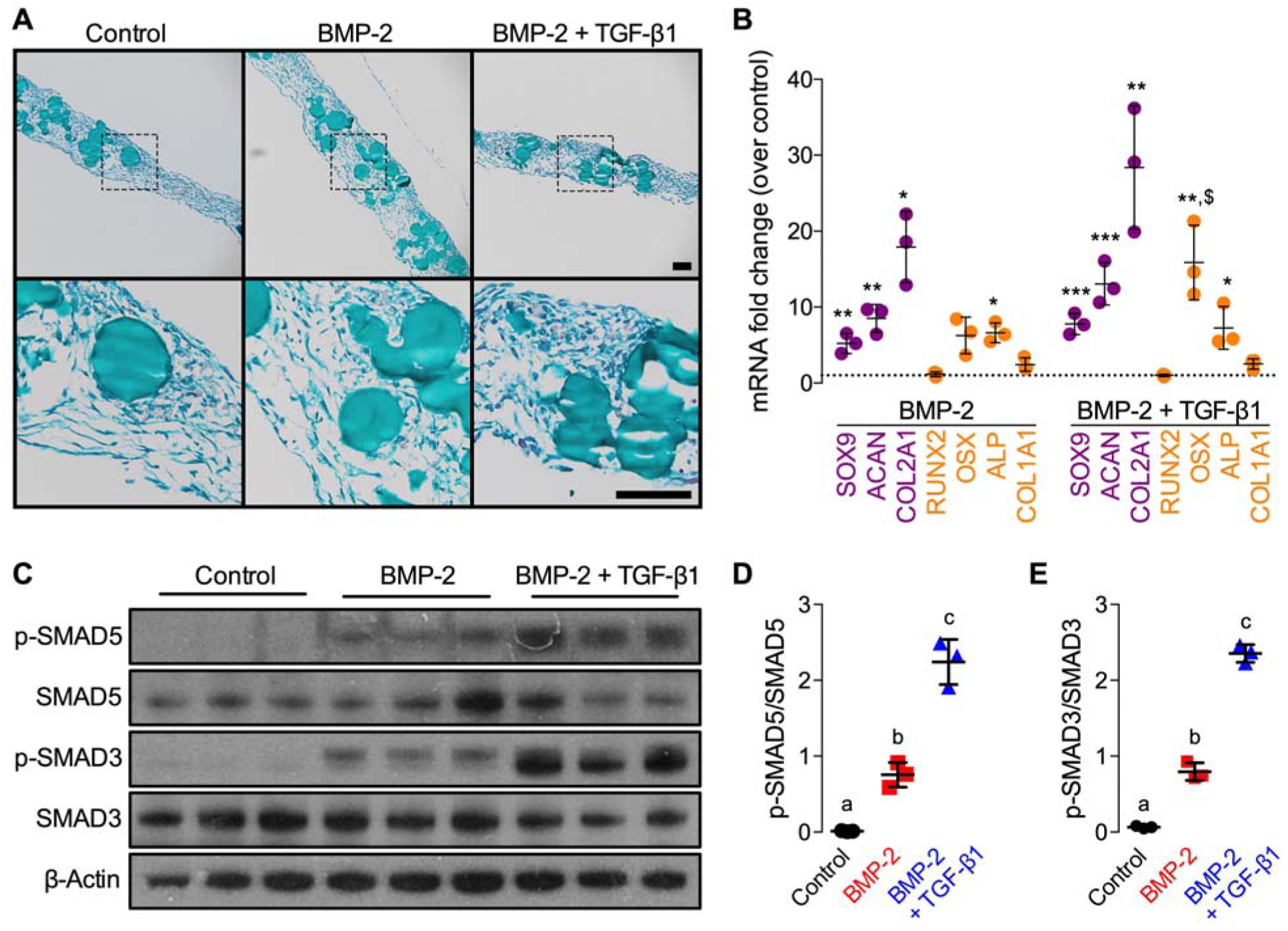
Effects of morphogen priming of engineered hMSC condensations on *in vitro* chondrogenic lineage specification at the time of implantation. Histological Safranin-O/Fast green staining of representative hMSC condensations at the time of implantation (2 days; N = 3 per group). Scale bars, 100 μm (10x: top; 40x: bottom, magnification of dotted squares). (**B**) Normalized mRNA fold-change over control of key chondrogenic or osteogenic markers by qRT-PCR (N = 3 per group; *p<0.05, **p<0.01, ***p<0.001 vs. control; ^$^p<0.05 vs. BMP-2-containing hMSC condensations). (**C**) Representative immunoblots and (**D**) relative quantification of p-SMAD5/SMAD5 and (**E**) p-SMAD3/SMAD3 in lysates of day 2 hMSC condensations (N = 3 per group). β-Actin served as loading control. Individual data points shown with mean ± SD. Comparisons between groups were evaluated by one-way ANOVA with Tukey’s post-hoc tests. Repeated significance indicator letters (a,b,c) signify p > 0.05, while groups with distinct indicators signify p < 0.05.

### In vivo tissue differentiation and composition

Next, to test the combinatorial effects of morphogen presentation and mechanical loading on local tissue differentiation, endochondral lineage progression, and matrix organization *in vivo*, we performed histological analyses of defect tissues at weeks 4 and 12. Control condensations exhibited predominantly fibrous and adipose tissue spanning the defects, and bone formation was apparent only capping the diaphyseal ends (Figure 5A,B; Supplementary Figures S7,8A,D,G,J). BMP-2-containing hMSC condensations induced the formation of primary woven bone and lamellar bone with lacunae-embedded osteoctyes surrounded by marrow-like tissue by week 4, and increased lamellar bone by week 12 (Fig. 5A). Aside from stiff control at 4 weeks, control and BMP-2-containing groups exhibited minimal Safranin-O-stained glycosaminoglycan (GAG) matrix at both time points. (Figure 5A,B; Supplementary Figures S7,8B,E,H,K). Co-delivery of BMP-2 + TGF-β1 induced robust bone formation exhibiting lacunae-embedded osteoctyes in well-defined trabeculae with peripheral positive GAG-staining, evidence of prior cartilaginous template transformation. Mechanical loading of BMP-2 + TGF-β1-containing condensations promoted formation of growth plate-like, transverse cartilage bands that featured zonal organization of mature and hypertrophic chondrocytes with prominent Safranin-O-stained GAG matrix embedded in trabecular bone and aligned orthogonal to the principal ambulatory load axis (Figure 5A,B; Supplementary Figures S7,8C,F,I,L). Hypertrophic chondrocytes and new bone formation at the interface were indicative of active endochondral bone formation in the dual morphogen group with mechanical loading at both 4 and 12 weeks (Fig. 5B).

Together, these data suggest that, though both BMP-2 and BMP-2 + TGF-β1 induced chondrogenic priming prior to implantation, endochondral ossification *in vivo* was apparent only in groups with TGF-β1 co-presentation. Further, *in vivo* mechanical cues potentiated cartilage formation and prolonged endochondral ossification.

**Figure 4:**
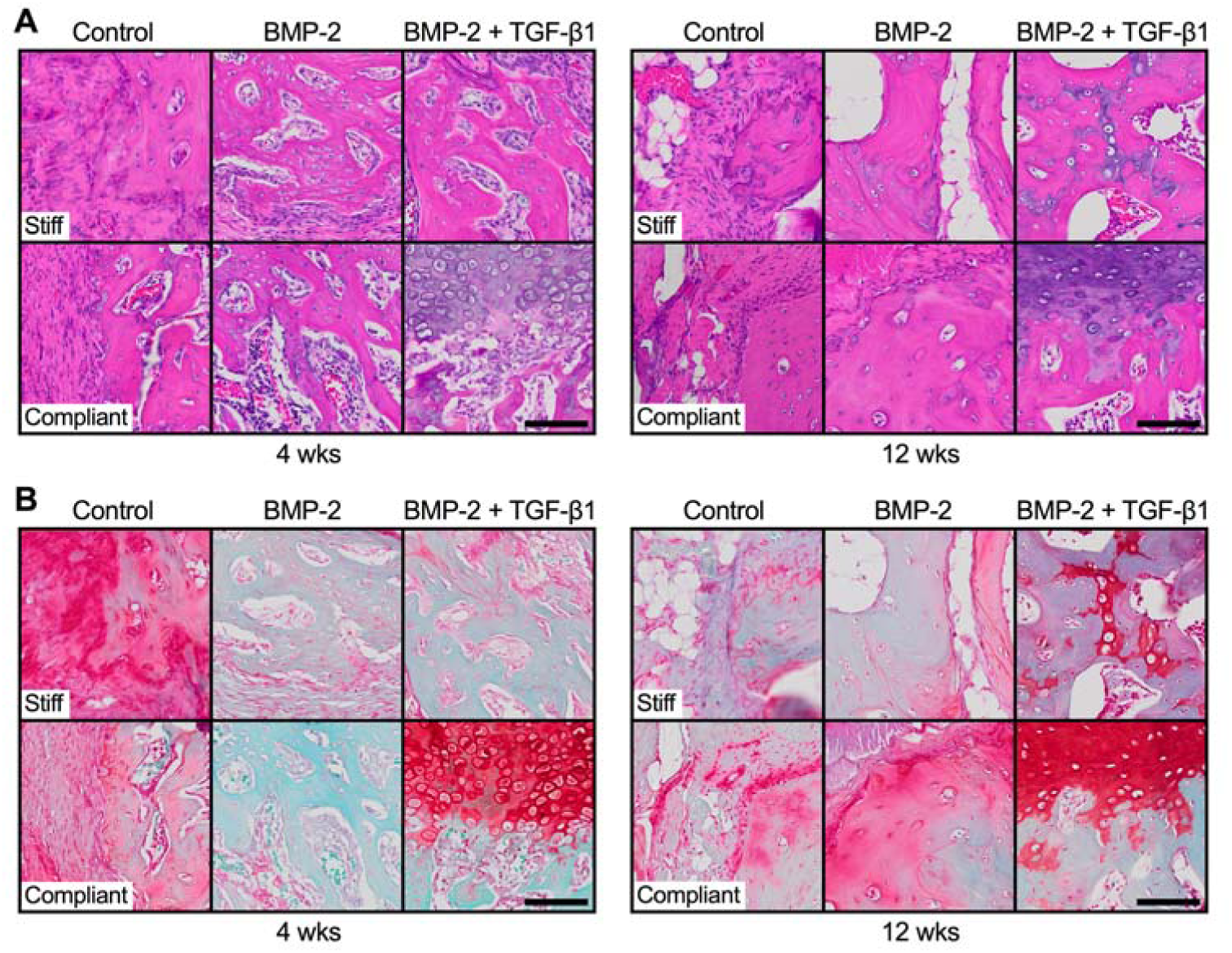
Effects of morphogen priming of engineered hMSC condensations and *in vivo* mechanical loading on tissue-level bone regeneration. Representative histological (**A**) H&E and (**B**) Safranin-O/Fast green staining of defect tissue at week 4 (left) and week 12 (right), with stiff (top) and compliant fixation (bottom), selected based on mean bone volume. Scale bars, 100 μm (40x).

## Discussion

The aim of this study was to replicate the cellular, biochemical, and mechanical environment present during limb development for functional regeneration of large segmental bone defects. Specifically, we used (i) engineered mesenchymal condensations formed by cellular self-assembly, which contained (ii) microparticle-mediated growth factor presentation to activate specific morphogenetic pathways *in situ*, and which, upon implantation, were exposed to (iii) *in vivo* mechanical loading. We tested the hypothesis that TGF-β1 and/or BMP-2 presentation from encapsulated microparticles within engineered hMSC condensations would promote endochondral regeneration of critical-sized femoral defects in a manner dependent on the *in vivo* mechanical environment. While both BMP-2 and BMP-2 + TGF-β1 presentation induced chondrogenic priming at the time of *in vivo* transplantation, endochondral ossification was apparent only in the dual morphogen group and was enhanced by mechanical loading. Specifically, *in vivo* ambulatory mechanical loading significantly enhanced the rate of bone formation in the four weeks after load initiation in the dual morphogen group, improved bone distribution in the callus, and fully restored mechanical bone function. In contrast, mechanical loading had no effect on bone regeneration in control groups without morphogen presentation, and likewise had no effect on autograft-mediated repair.

Multiple reports have described self-assembled hMSC condensations to form cartilage templates that can undergo hypertrophy and progress through endochondral ossification *in vivo* (*25, 26, 29, 34-37*). In these studies, chondrogenic priming was achieved by means of exogenously supplied morphogens, requiring *in vitro* pre-culture of 3 weeks and longer (*25, 26, 29, 34-37*). This requires time and associated costs and limits the precision of morphogen spatial distribution control. Further, few studies to date have achieved function-restoring bone formation in orthotopic models using this strategy (*28, 29, 54*). We previously demonstrated that sequential *in situ* morphogen presentation to hMSC condensations, utilizing mineral-coated hydroxyapatite microparticles (*49*) and crosslinked gelatin microspheres (*39*) to control the bioavailability of BMP-2 and TGF-β1, respectively, facilitates both chondrogenic and osteogenic differentiation *in vitro* (*41*), and promotes calvarial bone regeneration via endochondral ossification *in vivo* (*27*). Here, we show that this spatiotemporally controllable and localized morphogen delivery strategy, inspired by early limb development, eliminates the need for time- and cost-ineffective pre-differentiation of the cellular constructs and achieved mechanically-functional regeneration without the ectopic bone formation associated with BMP-2/collagen.

In addition to their efficacy in morphogen presentation, the condensations facilitated endochondral healing by providing a non-structural, immature intermediate, much like a callus in fracture healing or cartilage anlage in limb development. We showed previously that structural scaffolds that mimic the material properties of mature bone shield tissue from the stimulatory and beneficial effects of mechanical load during healing (*22*), suggesting that having a flexible intermediate structure is more suitable graft material for mechanical regulation of bone regeneration. Here, we found that *in vivo* mechanical loading via compliant fixation exerted stimulatory effects on defect healing, particularly in the period of plate actuation (4-8 wks), which resulted in complete functional bone regeneration, i.e., restoration of biomechanical competency comparable to un-operated, intact controls. These data indicate the importance of morphogen presentation for stem cell-mediated regeneration of bone defects, and potentially imply that the high stiffness of autograft bone may interfere with load-induced bone formation.

Compressive interfragmentary motion is necessary for cartilaginous callus formation and endochondral ossification during fracture healing (*9, 55*), and here the presence of growth plate-like cartilage structures, exhibiting zonal organization of mature and hypertrophic chondrocytes embedded in marrow-containing trabecular bone, suggests that BMP-2 + TGF-β1-containing hMSC condensations facilitated defect healing chiefly through endochondral ossification. This was consistent with a recent study demonstrating that a chondrogenic response to BMP-4 is dependent upon transient activation of TGF-β signaling in the early limb bud (*53*). *In vitro* analysis confirmed robust chondrogenic priming of the cellular constructs at the time of surgery. While this was also the case with BMP-2-only presenting condensations, upon defect implantation these constructs stimulated overall inferior bone regeneration compared to dual morphogen groups independent of the *in vivo* mechanical environment, and in a manner akin to intramembranous ossification that was consistent with previous reports (*44, 45, 47*). Nevertheless, no ectopic bone formation, as seen with BMP-2 soaked on collagen at ∼2 µg (*56*), was observed, similar to autografts as the other clinical standard we initially tested our technology against. This suggests an improved safety profile in the context of BMP-2 delivery from scaffold-free, self-assembled cellular constructs.

In conclusion, this study presents a human progenitor cell-based bone tissue engineering approach that recapitulates certain aspects of the normal endochondral cascade in early limb development. Implantation of chondrogenically-primed high-density cellular condensations, achieved through *in situ* morphogen presentation rather than lengthy pre-culture, in large bone defects that would otherwise not heal if left untreated, re-established biomechanical competency in limbs stabilized with custom compliant fixation plates with elective actuation at 4 weeks, after stable fixation to initiate bone regeneration. Further studies may elucidate the role of elective actuation timing in this regenerative strategy. Our findings are of clinical relevance and advance the current understanding in the growing field of developmental engineering. Furthermore, the system described herein can be used to study the complex biophysical mechanisms that govern tissue regeneration in health and disease.

## Materials and Methods

### Study design

The objective of this work was to mimic the cellular, biochemical, and mechanical environment of the endochondral ossification process during early limb development via *in situ* morphogen priming of high-density hMSC condensations and controlled *in vivo* mechanical cues upon implantation in large bone defects. We employed the established critical-size rat femoral segmental defect model in 12-week-old male Rowett nude (RNU) rats with custom internal fixation plates that allow controlled transfer of ambulatory loads *in vivo.* The sample size was determined with G*Power software (*57*) based on a power analysis using population standard deviations and estimated effect sizes from our prior studies (*40, 45*). The power analysis assumed a two-tailed alpha of 0.05, power of 0.8, and effect sizes of ranging from 0.1 to 0.3. A minimum sample number of N = 6 per group was computed, with an ideal sample number of N = 12 for all non-destructive and destructive analyses per time point. An N = 10 was selected for all non-destructive and destructive analyses per time point, accommodating a 5-10% complication rate consistent with our prior studies. Animals were randomly assigned to the treatment groups for both studies. Where indicated, limbs were excluded from the analysis based on radiographic evidence of fixation plate failure. Data collection occurred at predetermined time points informed by previous studies. All analyses were performed by examiners blinded to the treatment group.

### Experiments and study groups

**<u>Initial studies</u>**: For the first initial study, the experimental design featured one treatment group with two mechanical loading conditions. Defects received morselized autograft autograft prepared by mincing the excised cortical biopsy in sterile phosphate buffered saline (PBS) **[Autograft]** contained within an electrospun, perforated PCL nanofiber mesh tube. Limbs were stabilized with stiff **[Stiff]** or axially-compliant **[Compliant]** fixation plates initially implanted in a locked configuration to prevent loading (k_axial_ = 250 ± 35 N/mm), but after four weeks the plates were surgically unlocked to enable load transfer (k_axial_ =8.0 ± 3.5 N/mm) (N = 6-8 per group) (*43-45*). For the second initial study, the experimental design featured three treatment groups with one mechanical loading condition. Defects received 1) three hMSC condensations for a total of 6.0×10^6^ cells with bone morphogenetic protein-2 (BMP-2)-loaded mineral-coated hydroxyapatite (MCM) microparticles (1.9 µg) and unloaded gelatin microspheres (GM) **[BMP-2 (hMSCs)]**, 2) BMP-2 (1.9 µg) soaked onto 8 mm pre-cut absorbable collagen sponge (INFUSE(tm) Bone Graft; Medtronic, Memphis, TN) 15 min prior to implantation **[BMP-2 (collagen)]**, or 3) morselized autograft in sterile PBS **[Autograft]**, each contained within an electrospun, perforated PCL nanofiber mesh tube. Limbs were stabilized with stiff fixation plates **[Stiff]** that limit load transfer (k_axial_ = 260 ± 28 N/mm) (N = 7-10 per group), modified from prior studies (*43-45*). **<u>Main study:</u>** The experimental design featured three treatment groups with two mechanical loading conditions (Table 1). Defects received three hMSC condensations for a total of 6.0×10^6^ cells incorporated with 1) unloaded MCM and GM **[Control]**, 2) BMP-2-loaded MCM (1.9 µg) and unloaded GM **[BMP-2]**, or 3) BMP-2-loaded MCM (1.9 µg) and transforming growth factor-β1 (TGF-β1)-loaded GM (1.8 µg) **[BMP-2+TGF-**β**1]** contained within an electrospun, perforated PCL nanofiber mesh tube. Limbs were stabilized with stiff **[Stiff]** or axially-compliant **[Compliant]** fixation plates initially implanted in a locked configuration to prevent loading (k_axial_ = 250 ± 35 N/mm), but after four weeks the plates were surgically unlocked to enable load transfer (k_axial_ = 8.0 ± 3.5 N/mm) (N = 3-10 per group) (*43-45*).

### hMSC isolation and expansion

Human mesenchymal stromal/stem cells (hMSCs) were derived from the posterior iliac crest of two healthy male donors (26 and 41 years of age) using a protocol approved by the University Hospitals of Cleveland Institutional Review Board. Cells were isolated using a Percoll gradient (Sigma-Aldrich, St. Louis, MO) and cultured in low-glucose Dulbecco’s modified Eagle’s medium (DMEM-LG; Sigma-Aldrich, St. Louis, MO) containing 10% fetal bovine serum (FBS; Sigma-Aldrich), 1% penicillin/streptomycin (P/S; Fisher Scientific), and 10 ng/ml fibroblast growth factor-2 (FGF-2, R&D Systems, Minneapolis, MN) (*39, 41, 48, 58*).

### Hydroxyapatite microparticle mineral coating and BMP-2 loading

MCM were kindly provided by Dr. William L. Murphy (University of Wisconsin, Madison, WI). Their preparation using low carbonate (4.2 mM NaHCO_3_) coating buffer and detailed characterization has been reported previously (*41*). Lyophilized MCM from the same batch as used in these prior studies, and our own (*48*), were loaded with a 100 µg/ml solution of recombinant human (rh) BMP-2 (Dr. Walter Sebald, Department of Developmental Biology, University of Würzburg, Germany; 6.4 µg/mg) in PBS for 4 h at 37°C. BMP-2-loaded MCM were then centrifuged at 800xg for 2 min and washed 2x with PBS. Unloaded MCM without growth factor were incubated with PBS only and treated similarly.

### Gelatin microsphere synthesis and TGF-β1 loading

Gelatin microspheres (GM) (*41, 48, 59,60*) were synthesized from 11.1% (w/v) gelatin type A (Sigma-Aldrich) using a water-in-oil single emulsion technique and crosslinked for 4 h with 1% (w/v) genipin (Wako USA, Richmond, VA) (60). Hydrated GM were light blue in color and spherical in shape with an average diameter of 52.9±40.2 μm and a crosslinking density of 25.5 ± 7.0% (*48*). Growth factor-loaded microspheres were prepared by soaking crosslinked, UV-sterilized GM in 80 μg/ml solution of rhTGF-β1 (Peprotech, Rocky Hill, NJ) in PBS for 2 h at 37°C. Unloaded microspheres without growth factor were hydrated similarly using only PBS.

### Preparation of microparticle-incorporated mesenchymal condensations

Expanded hMSCs (2.0×10^6^ cells/construct; passage 4) were thoroughly mixed with BMP-2-loaded MCM (1.6 µg/mg; 0.4 mg/sheet) and TGF-β1-loaded GM (0.4 µg/mg; 1.5 mg/construct) in chemically defined medium [DMEM-HG (Sigma-Aldrich), 1% ITS+ Premix (Corning), 1 mM sodium pyruvate (HyClone), 100 μM non-essential amino acids (Lonza), 100 nM dexamethasone (MP Biomedicals, Solon, OH), 0.05 mM L-ascorbic acid-2-phosphate (Wako), and 1% P/S (Fisher Scientific)] (*27, 39*). Five hundred microliter of the suspension were seeded onto the pre-wetted membrane of transwell inserts (3 µm pore size, 12 mm diameter; Corning) and allowed to self-assemble for 2 days. After 24 h, medium in the lower compartment was replaced. Control constructs containing either unloaded MCM and/or GM were prepared and cultured in a similar fashion. After 48 h, hMSC condensations were harvested for implantation.

### Nanofiber mesh production

Nanofiber meshes were formed by dissolving 12% (w/v) poly-(ε-caprolactone) (PCL; Sigma-Aldrich) in 90/10 (v/v) hexafluoro-2-propanol/dimethylformamide (Sigma-Aldrich). The solution was electrospun at a rate of 0.75 ml/h onto a static aluminum collector. 9 mm x 20 mm sheets were cut from the product, perforated with a 1 mm biopsy punch (VWR, Radnor, PA), and glued into tubes around a 4.5 mm mandrel with UV glue (Dymax, Torrington, CT). Meshes were sterilized by 100% ethanol evaporation under UV light over-night and washed 3x with sterile PBS before implantation.

### Surgical procedure

Critical-sized (8 mm) bilateral segmental defects were created in the femora of 12-week-old male Rowett nude (RNU) rats (Taconic Biosciences Inc., Hudson, NY) under isoflurane anesthesia (*61*). Limbs were stabilized by custom internal fixation plates that allow controlled transfer of ambulatory loads in vivo (*43*) and secured to the femur by four bi-cortical miniature screws (J.I. Morris Co, Southbridge, MA). Animals were given subcutaneous injections of 0.04 mg/kg buprenorphine every 8 h for the first 48 h postoperative (post-op) and 0.013 mg/kg every 8 h for the following 24 h, with or without 4-5 mg/kg carprofen every 24 h for 72 h. In addition, 5 ml of 0.9% NaCl were administered subcutaneously to aid in recovery. All procedures were performed in strict accordance with the NIH Guide for the Care and Use of Laboratory Animals, and the policies of the Institutional Animal Care and Use Committee (IACUC) at Case Western Reserve University (Protocol No. 2015-0081) and the University of Notre Dame (Protocol No. 14-05-1778).

### In vivo X-ray and microCT

*In vivo* X-rays were obtained using an Xtreme scanner (Bruker, Billerica, MA) at 45 kVp, 0.4 mA, and 2 s exposure time. A binary bridging score was assigned by two independent, blinded observers, and determined as mineralized tissue fully traversing the defect. *In vivo* microCT scans were performed at or 4, 8, and 12 weeks to assess longitudinal defect healing. For initial studies, animals were scanned using an Inveon microPET/CT system (Siemens Medical Solutions, Malvern, PA) at 45 kVp, 0.2 mA, and 35 μm isotropic voxels. Data were reconstructed using system-default parameters for analyzing bone and accounting for the metal in the fixation plates. DICOM-exported files were processed for 3-D analysis (CTAn software, Skyscan; Bruker) using a gauss filter at 1.0 pixel radius and a global threshold range of 28-255 for all samples. Bone volume was determined in a standard region of interest (ROI) spanning the length of the defect. For the main study, animals were scanned using an Albira PET/SPECT/CT system (Bruker) at 45 kVp, 0.4 mA, and 125 μm voxel size. A global threshold was applied for each data set, and bone volume determined in a standard ROI spanning the length of the defect.

### Ex vivo microCT

After 12 weeks, the animals were euthanized by CO_2_ asphyxiation and hind limbs were excised for high resolution microCT analysis. Data were acquired using a Skyscan 1172 microCT scanner (Bruker) with a 0.5 mm aluminum filter at 75 kVp and 0.1 mA. Femora wrapped in gauze were placed in a plastic sample holder with the long axis oriented parallel to the image plane, and scanned in PBS at 20 µm isotropic voxels, 560 ms integration time, rotation step of 0.5°, and frame averaging of 5. All samples were scanned within the same container using the same scanning parameters. All scans were then reconstructed using NRecon software (Skyscan) with the same reconstruction parameters (ring artifact reduction of 5, beam hardening correction of 20%). For 3-D analysis (CTAn software, Skyscan), a gauss filter at 1.0 pixel radius and a global threshold range of 65-255 were used. This segmentation approach allowed viewing of the normal bone architecture in the binary images as seen in the original reconstructed images (*62*). Three hundred and twenty-five slices in the center of each defect were analyzed in a standard ROI using a 10 mm (total) or 5 mm (defect) diameter circle centered on the medullary canal. Bone volume, bone volume fraction, polar moment of inertia (pMOI), and the morphometric parameters trabecular number, trabecular thickness, trabecular separation, structure model index, degree of anisotropy, and connectivity density were calculated. Trabecular morphometry and pMOI of three age-matched femora were analyzed in the same manner for comparison. Proximal and distal total bone volume were calculated by halving the slice number in each sample and separately segmenting each half for comparison. All representative images were chosen based on average bone volume values.

### Biomechanical testing

Femora excised at 12 weeks were biomechanically tested in torsion to failure. Limbs were cleaned of soft tissue and fixation plates were carefully removed. Bone ends were potted in Wood’s metal (Alfa Aesar; Fisher Scientific), mounted on a Mark-10 TSTM-DC test stand with MR50-12 torque sensor (1.35 N-m) and 7i torque indicator (Mark-10 Corp., Copiague, NY) using custom fixtures, and tested to failure at a rate of 3°/s. For each sample, maximum torque at failure was recorded and torsional stiffness computed as the slope of the linear region in the torque-rotation curve. Samples were compared to 3 age matched, un-operated femurs.

### Histological analysis

hMSC condensations (N = 3/group) were fixed in 10% neutral buffered formalin (NBF) for 24 h at 4°C before switching to 70% ethanol. One representative femur per group was taken for histology at weeks 4 and 12 post-surgery, chosen based on microCT-calculated average bone volumes. Femora were fixed in 10% NBF for 72 h at 4°C and then transferred to 0.25 M ethylenediaminetetraacetic acid (EDTA) pH 7.4 for 14 d at 4°C under mild agitation on a rocker plate, with changes of the decalcification solution every 3-4 days. Following paraffin processing, 5 µm mid-sagittal sections were cut using a microtome (Leica Microsystems Inc., Buffalo Grove, IL) and stained with hematoxylin & eosin (H&E) and Safranin-O/Fast-green (Saf-O). Light microscopy images were captured using an Olympus BX61VS microscope (Olympus, Center Valley, PA) with a Pike F-505 camera (Allied Vision Technologies, Stadtroda, Germany).

### Quantitative reverse transcription-polymerase chain reaction (qRT-PCR) analysis

hMSC condensations (N = 3/group) were homogenized in TRI Reagent (Sigma-Aldrich) for subsequent total RNA extraction and cDNA synthesis (iScript™ kit; Bio-Rad, Hercules, CA). One hundred nanograms of cDNA were amplified in duplicates in each 40-cycle reaction using a Mastercycler (Eppendorf, Hauppauge, NY) with annealing temperature set at 60°C, SYBR® Premix Ex Taq™ II (Takara Bio Inc., Kusatsu, Shiga, Japan), and custom-designed qRT-PCR primers (Table S1; Life Technologies, Grand Island, NY). Transcript levels were normalized to GAPDH and gene expression was calculated as fold-change using the comparative C_T_ method (*63*).

### Immunoblotting

hMSC condensations (N = 3/group) were homogenized in CelLytic™ MT lysis buffer (Sigma-Aldrich) supplemented with Halt™ protease and phosphatase inhibitor cocktail (Thermo Scientific). Equal amounts (15 µg) of protein lysates, determined by standard BCA protein assay kit (Pierce; Thermo Fisher Scientific), were subjected to SDS-PAGE using 10% NuPAGE® Bis-Tris gels (Invitrogen; Thermo Fisher Scientific) and transferred to 0.45 µm PVDF membranes (Millipore, Billerica, MA). Membranes were blocked with 5% bovine serum albumin (BSA) in standard TBST. The phosphorylation of intracellular SMAD3 and SMAD5 was detected using specific primary antibodies (anti-phospho-SMAD3 [ab52903]; anti-phospho-SMAD5 [ab92698]: Abcam, Cambridge, MA) followed by HRP-conjugated secondary antibodies (Jackson ImmunoResearch, West Grove, PA). Subsequently, the blots were stripped (Western Blot Stripping Buffer, Pierce; Thermo Fisher Scientific) and re-probed for the detection of the respective total protein (anti-SMAD3 [ab40854]; anti-SMAD5 [ab40771]: Abcam) and loading control (anti-β-Actin [A1978]: Sigma-Aldrich) with respective HRP-conjugated secondary antibodies (Jackson ImmunoResearch). Bound antibodies were visualized with the ECL detection system (Pierce; Thermo Fisher Scientific) on autoradiography film (Thermo Fisher Scientific). The intensity of immunoreactive bands was quantified using ImageJ software (National Institutes of Health, Bethesda, MD).

### Statistical analysis

Differences in bone volume and bone volume accumulation rate by *in vivo* microCT at weeks 4, 8, and 12 were determined by two-way analysis of variance (ANOVA) with Tukey’s *post hoc* test. Defect bridging was determined by chi-square test for trend in each group; comparisons between groups were assessed with individual chi-square tests and Bonferroni correction for multiple comparisons. *Ex vivo* microCT bone volume, bone volume fraction, and 3-D morphometry were assessed by one- or two-way ANOVA with interaction and Tukey’s *post hoc* test. Biomechanical properties were analyzed by two-way ANOVA with interaction and Tukey’s *post hoc* test. Mechanical property regressions were performed using an exhaustive best-subsets algorithm to determine the best predictors of maximum torque and torsional stiffness from a subset of morphologic parameters measured, including minimum or mean polar moment of inertia (J_min_ or J_mean_), bone volume fraction (BV/TV), binary bridging score (yes or no), trabecular thickness (Tb.Th), trabecular separation (Tb.Sp), trabecular number (Tb.N), degree of anisotropy (DA), and connectivity density (Conn.D) based on Akaike’s information criterion (AIC) (*51*). The lowest AIC selects the best model while giving preference to models with less parameters. Finally, the overall best model for each predicted mechanical property was compared to its measured value using type II general linear regression. All data are shown with mean ± SD, some with individual data points, or as box plots showing median as horizontal line, mean as +, and 25^th^ and 75^th^ percentiles as boxes with whiskers at minimum and maximum values. Fold-changes in mRNA expression and ratios of phosphorylated SMADs/total SMADs were analyzed by one-way ANOVA with Tukey’s multiple comparison *post hoc* test. The significance level was set at p<0.05 or lower. Groups with shared letters have no significant differences. GraphPad Prism software v6.0 (GraphPad Software, La Jolla, CA) was used for all analyses.

## Supplementary Materials

### Supplementary Figures

**Supplementary Figure S1:**
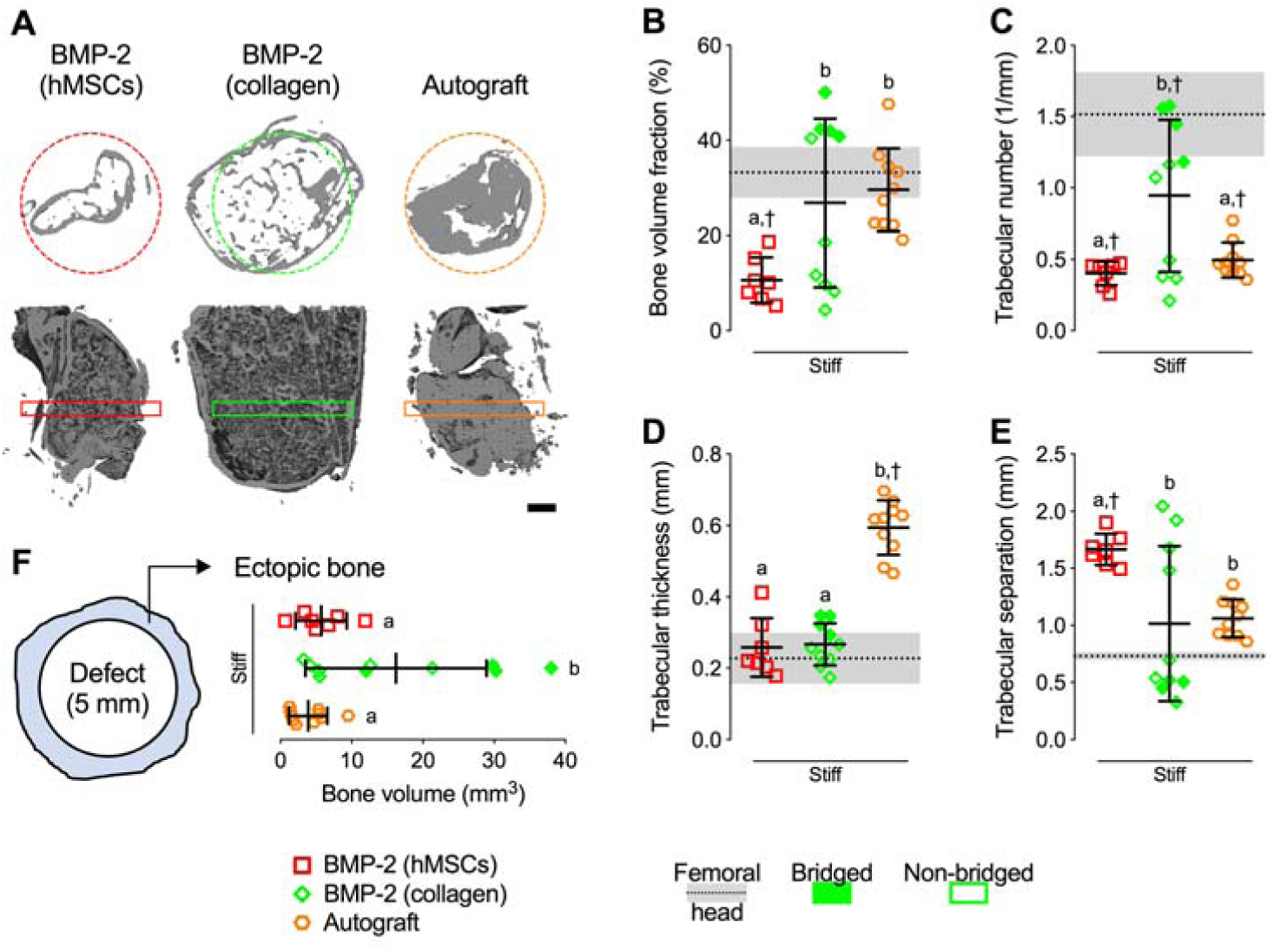
Effects of BMP-2-primed engineered hMSC condensations and routine clinical therapies on new bone quantity and architecture in absence of mechanical cues. (**A**) Representative 3-D microCT defect reconstructions of mid-shaft transverse (top) and sagittal (bottom) sections at week 12, selected based on mean bone volume. Dashed circles show 5 mm defect region. Rectangular boxes illustrate transverse cutting planes. Scale bar, 1 mm. (**B**) Morphometry analysis of bone volume fraction, (**C**) trabecular number, (**D**) trabecular thickness, (**E**) trabecular separation, shown with native femoral head properties (N = 3; dotted lines with gray shading: mean ± SD; ^†^p<0.05 vs. femoral head), and (**F**) ectopic bone formation (i.e., bone extending beyond the 5-mm defect diameter) at week 12 (N = 7-10 per group). Individual data points shown with mean ± SD. Comparisons between groups were evaluated by two-way ANOVA with Tukey’s post-hoc tests. Repeated significance indicator letters (a,b,c) signify p > 0.05, while groups with distinct indicators signify p < 0.05.

**Supplementary Figure S2:**
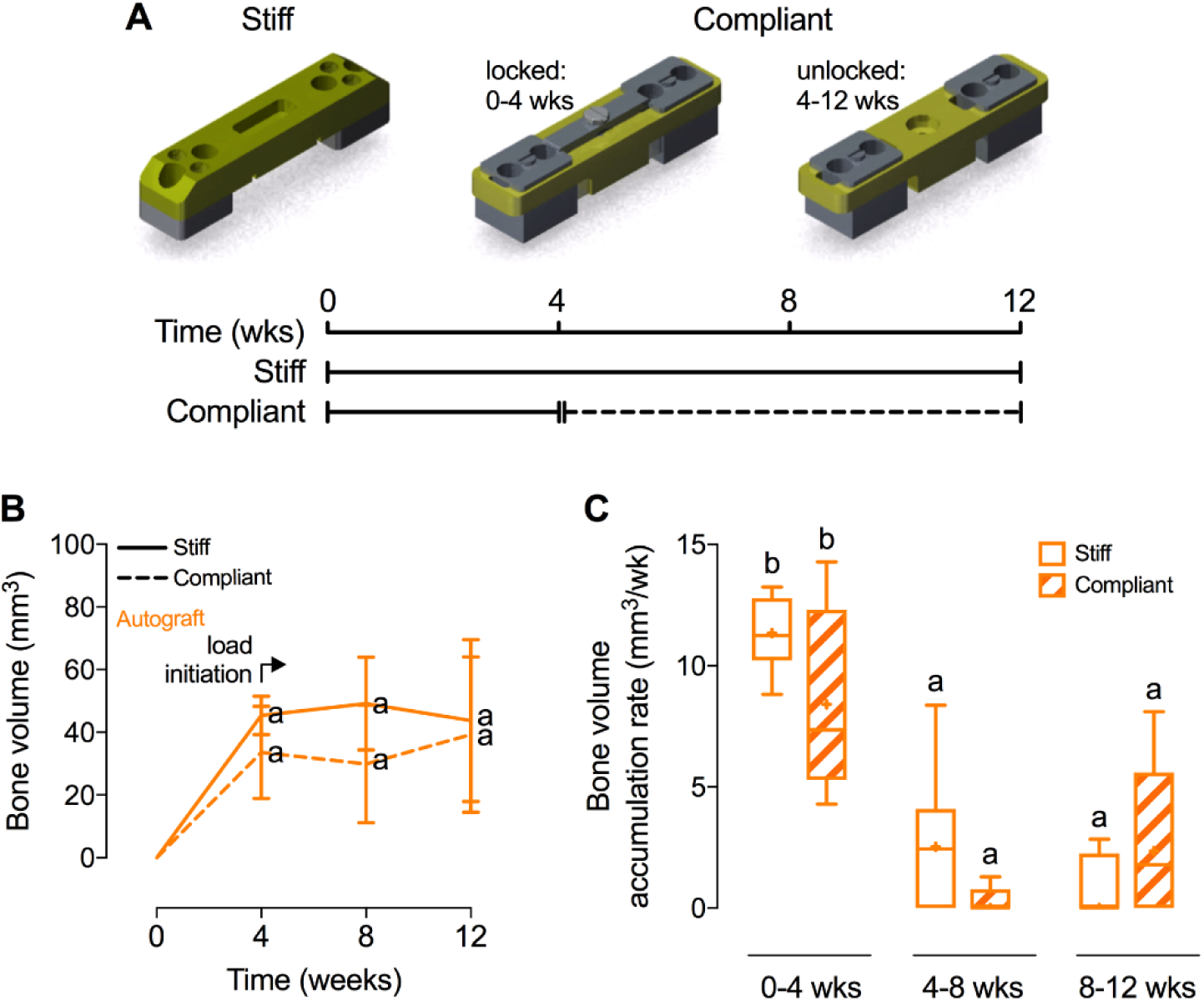
Effects of morselized autografts and *in vivo* mechanical loading on longitudinal bone formation and bone accumulation rate. (**A**) Stiff and compliant fixation plate configurations for dynamic control of ambulatory load transfer, and loading timeline with compliant plate unlocking at week 4. (**B**) Longitudinal quantification of bone volume at weeks 4, 8, and 12 by *in vivo* microCT (N = 6-8 per group). (**C**) Bone volume accumulation rate, defined as bone volume accrual over each 4-week interval. Data shown with mean ± SD. Box plots display median as horizontal line, mean as +, inter-quartile range as boxes, and min/max range as whiskers. Comparisons between groups were evaluated by two-way ANOVA with Tukey’s post-hoc tests. Repeated significance indicator letters (a,b,c) signify p > 0.05, while groups with distinct indicators signify p < 0.05 at each time point.

**Supplementary Figure S3:**
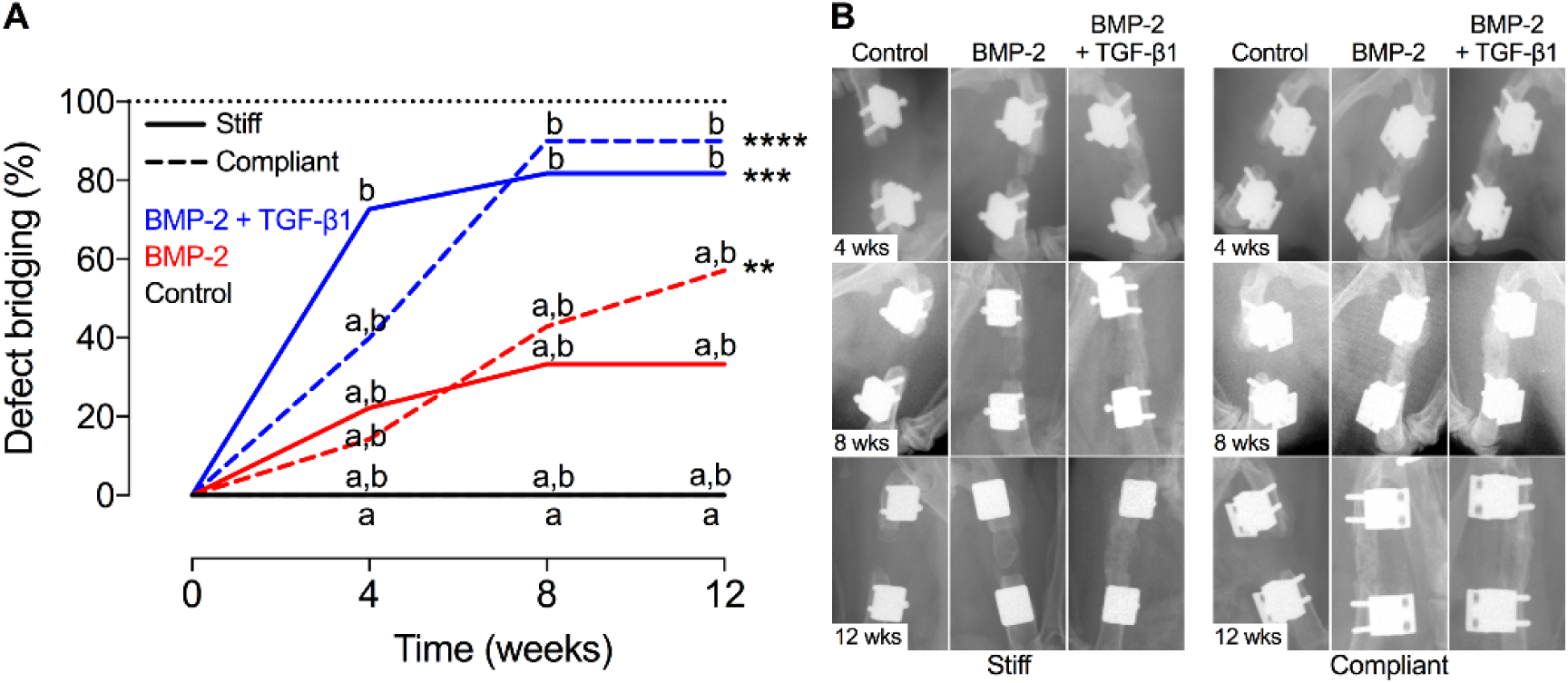
Effects of morphogen priming of engineered hMSC condensations and *in vivo* mechanical loading on defect bridging. (**A**) Longitudinal determination of defect bridging by *in vivo* radiography, defined as mineral fully traversing the defect (N = 4-11 per group). (**B**) Representative radiography images at 4, 8 and 12 weeks showing defect bridging per group over time, selected based on mean bone volume at week 12 (Figure 3). Significance of trend was analyzed by chi-square test (**p<0.01, ***p<0.001, ****p<0.0001). Differences between groups were determined by chi-square test at each time point with Bonferroni correction (p<0.01, correction factor of 5. Repeated significance indicator letters (a,b,c) signify p > 0.05, while groups with distinct indicators signify p < 0.05.

**Supplementary Figure S4:**
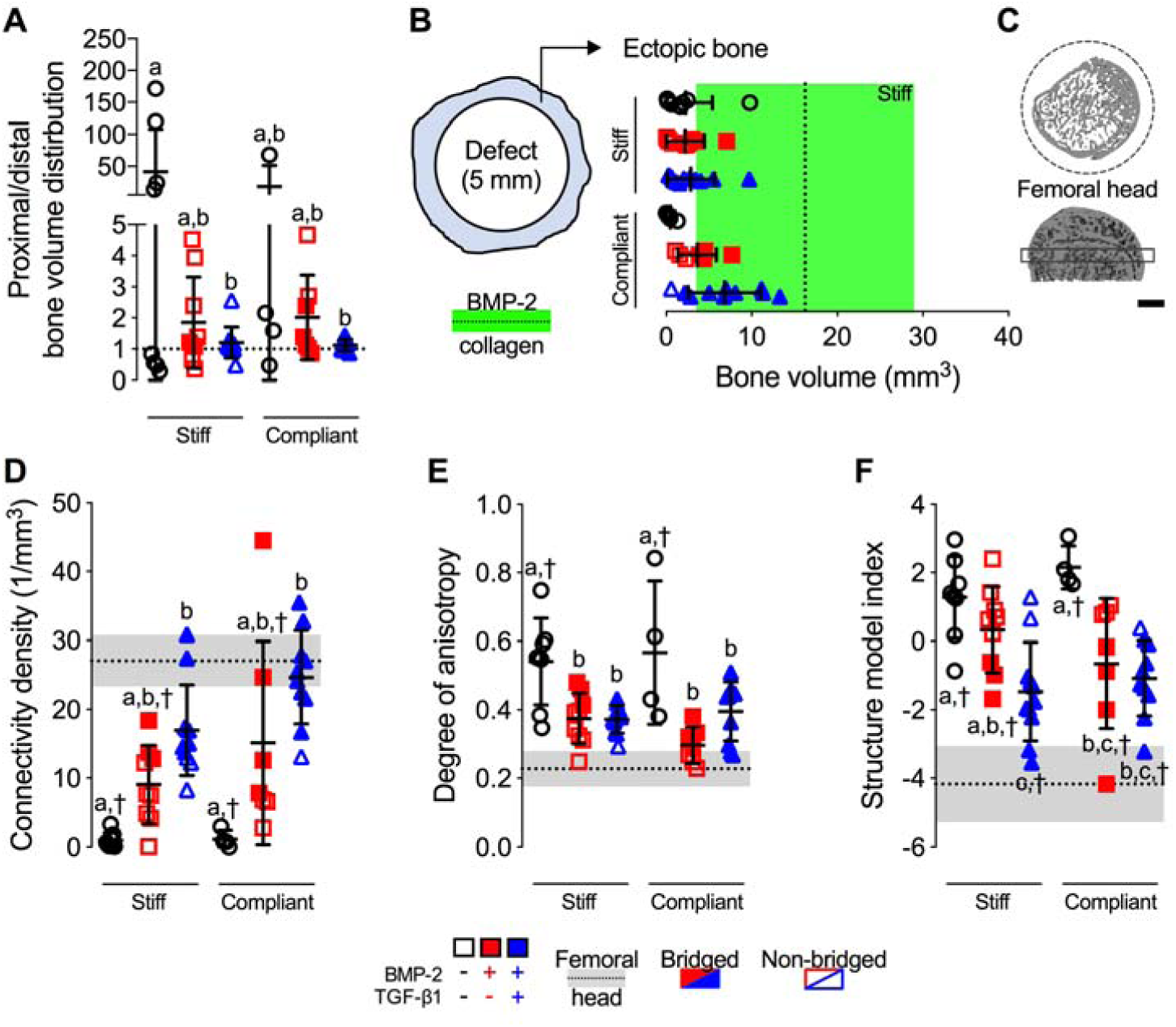
Effects of morphogen priming of engineered hMSC condensations and *in vivo* mechanical loading on new bone distribution and architecture. (**A**) Morphometry analysis of proximal vs. distal bone volume distribution (N = 4-11 per group; p<0.05), and (**B**) ectopic bone formation (i.e., bone extending beyond the 5-mm defect diameter) shown with BMP-2 soaked on collagen data (N = 9; dotted line with green shading: mean ± SD). (**C**) Representative 3-D microCT reconstructions of native femoral head transverse (top) and sagittal (bottom) sections, selected based on mean bone volume. Dashed circles show 5 mm defect region. Rectangular box illustrates transverse cutting plane. Scale bar, 1 mm. (**D**) Morphometry analysis of connectivity density, (**E**) degree of anisotropy, and (**F**) structure model index within the defect region (N = 4-11 per group), shown with native femoral head properties (N = 3; dotted lines with gray shading: mean ± SD; ^†^p<0.05 vs. femoral head). Individual data points shown with mean ± SD. Comparisons between groups were evaluated by two-way ANOVA with Tukey’s post-hoc tests. Repeated significance indicator letters (a,b,c) signify p > 0.05, while groups with distinct indicators signify p < 0.05.

**Supplementary Figure S5:**
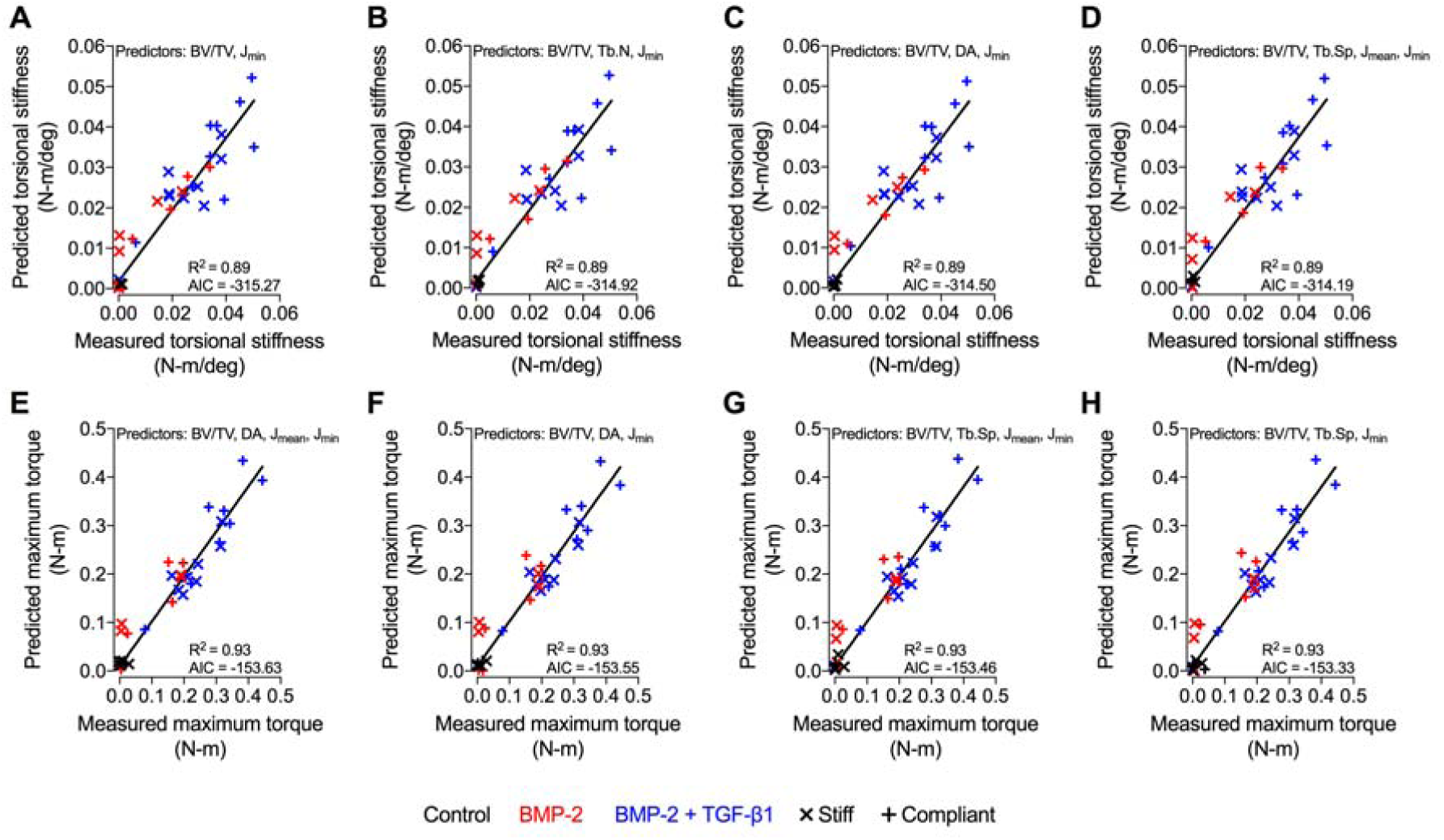
Best subsets analysis of mechanical testing data. Best subsets regression analysis of the top 5 models with selection made by minimization of Akaike’s Information Criterion (AIC). Predictive models of (**A-D**) torsional stiffness and (**E-H**) maximum torque at failure were composed of combinations of bone volume fraction (BV/TV), trabecular number (Tb.N), trabecular separation (Tb.Sp), minimum pMOI (J_min_), mean pMOI (J_mean_), and degree of anisotropy (DA). Type II regression was used to determine correlations (R^2^) between predicted and measured values.

**Supplementary Figure S6:**
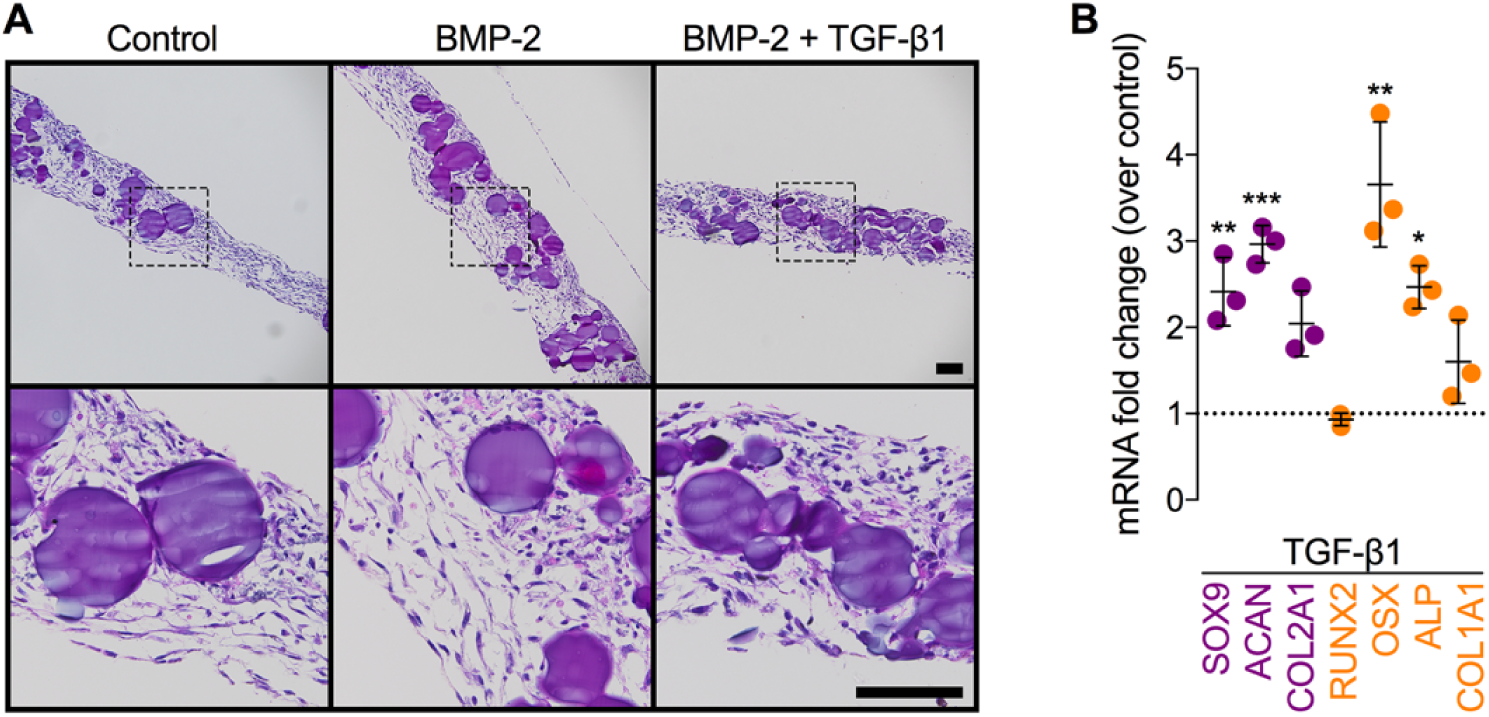
Effects of morphogen priming of engineered hMSC condensations on *in vitro* chondrogenic lineage specification at the time of implantation. (**A**) Histological H&E staining of representative hMSC condensations at the time of implantation (2 days; N = 3 per group). Scale bars, 100 μm (10x: top; 40x: bottom, magnification of dotted squares). (**B**) Normalized mRNA fold-change over control of key chondrogenic or osteogenic markers in TGF-β1-only loaded condensations by qRT-PCR (N = 3 per group; *p<0.05, **p<0.01, ***p<0.001 vs. control). Individual data points shown with mean ± SD. Analyzed by unpaired two-tailed Student’s *t* test.

**Supplementary Figure S7:**
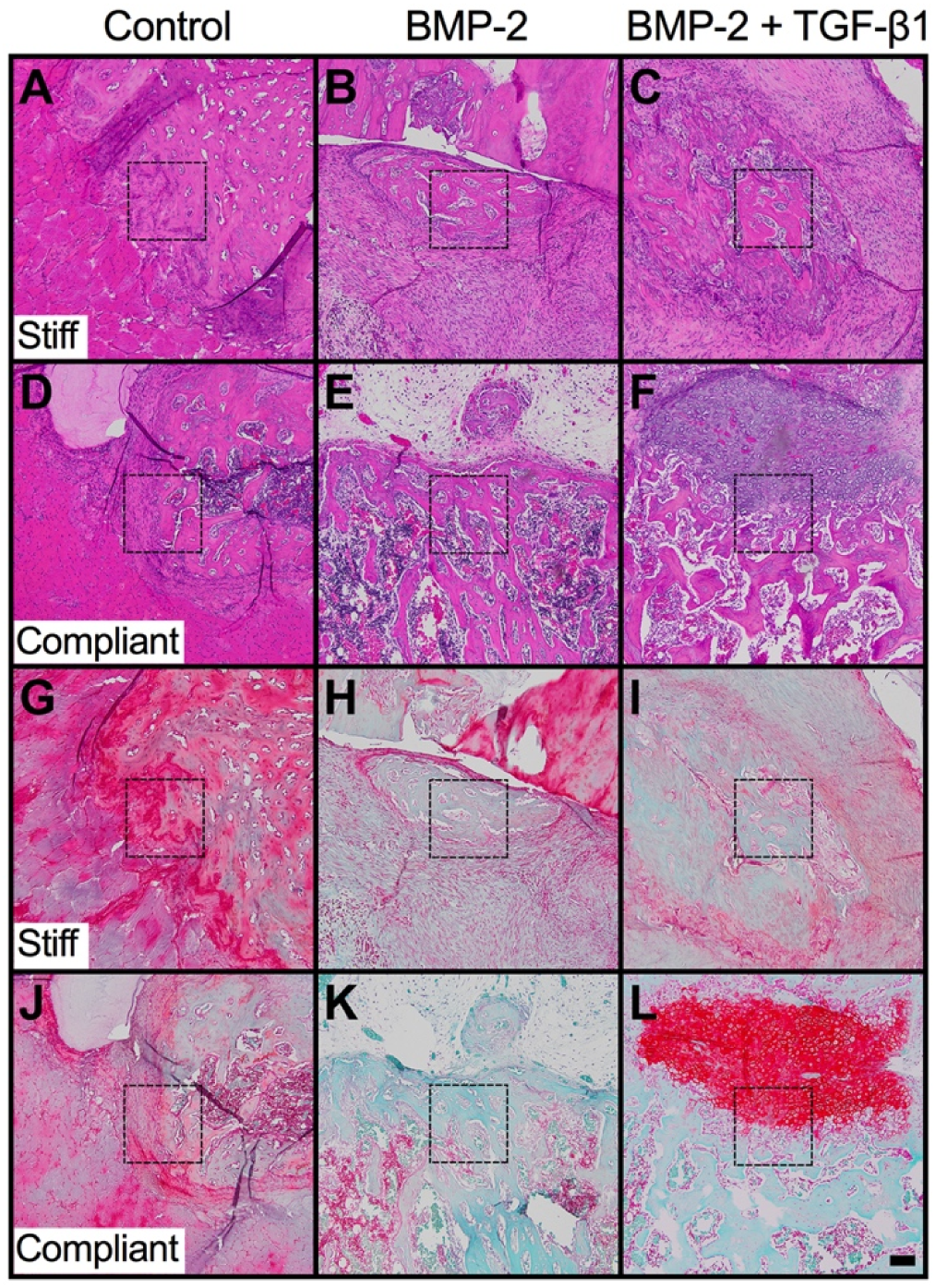
Effects of morphogen priming of engineered hMSC condensations and *in vivo* mechanical loading on tissue-level bone regeneration. Overview histology of bone formation at 4 weeks. (**A-F**) Representative histological H&E and (**G-L**) Safranin-O/Fast green staining of defect tissue at week 4, selected based on mean bone volume. Scale bar, 100 μm (10x; dotted squares show areas used in 40x images in Figure 4A,B).

**Supplementary Figure S8:**
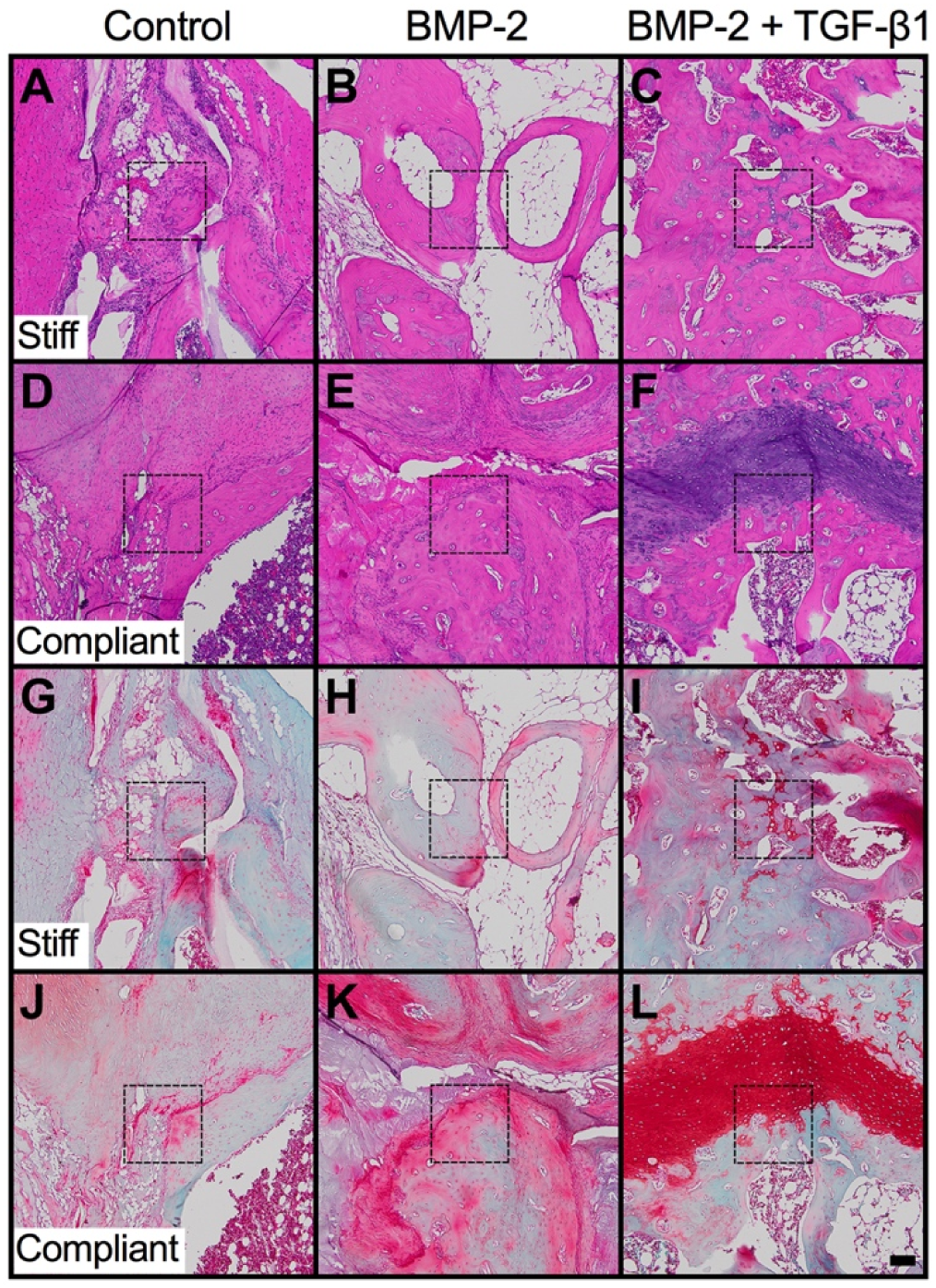
Effects of morphogen priming of engineered hMSC condensations and *in vivo* mechanical loading on tissue-level bone regeneration. Overview histology of bone formation at 12 weeks. **(A-F)** Representative histological H&E and (G-L) Safranin-O/Fast green staining of defect tissue at week 12, selected based on mean bone volume. Scale bar, 100 μm (10x; dotted squares show areas used in 40x images in Figure 4A,B).

### Supplementary Tables

**Table S1:**
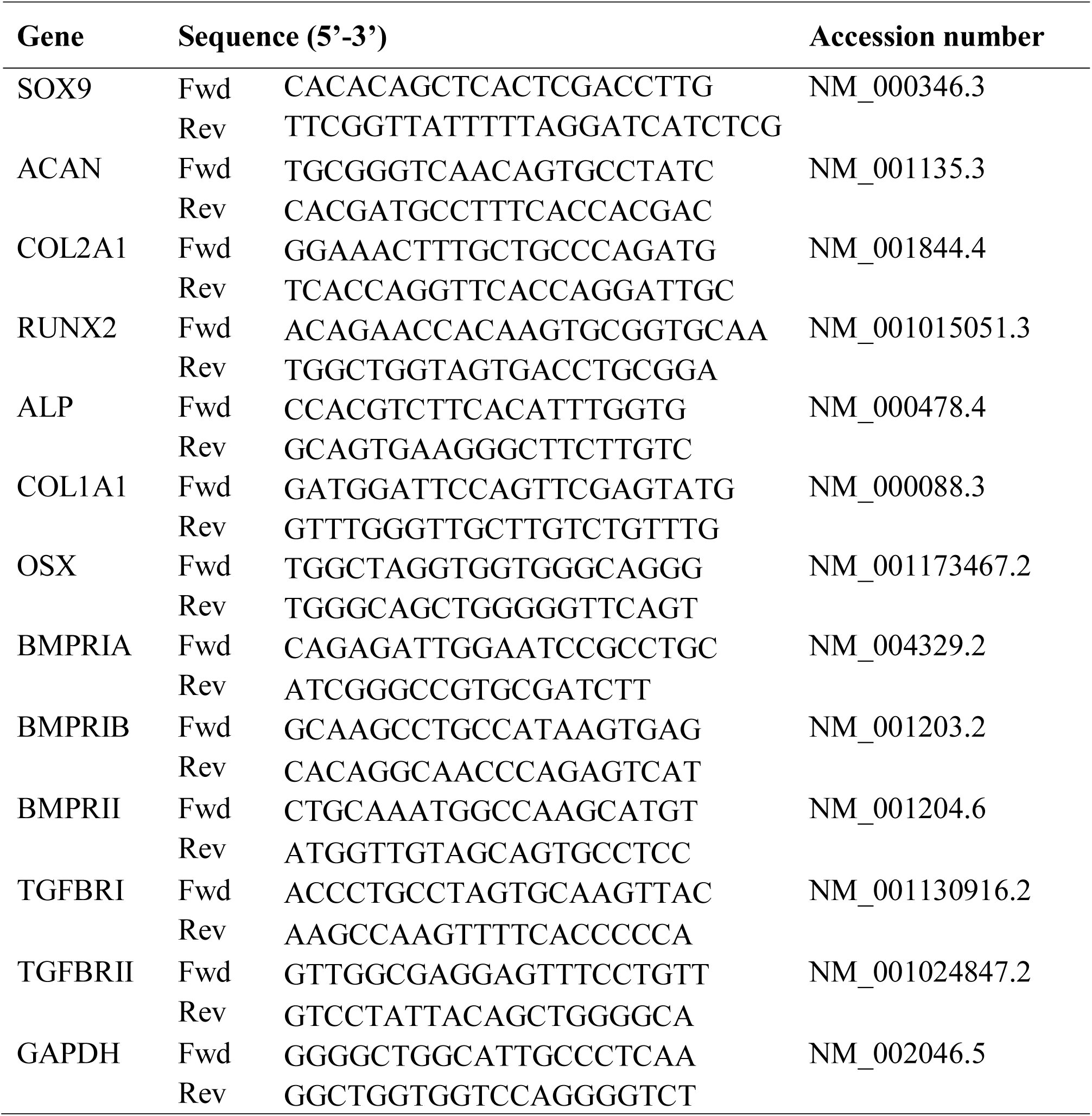
Oligonucleotide primer sequences for qRT-PCR.

## Acknowledgments

We thank the staff of the Freimann Life Science Center at the University of Notre Dame (ND), and the Animal Resource Center at Case Western Reserve University (CWRU) for animal care and husbandry. We also thank the staff of the ND Integrated Imaging Facility and the CWRU Imaging Research Core Facility for imaging support, Dr. Edward M. Greenfield at the CWRU Orthopaedic Research Facilities for imaging and biomechanical testing support, and Amad Awadallah at the CWRU Histology Core Facility for technical support. We also thank Dr. Xiaohua Yu and Dr. William L. Murphy at the University of Wisconsin, Madison, WI for providing the mineral-coated hydroxyapatite microparticles.

## Author contributions

S.H., A.M.M., R.T., J.D.B., and E.A. designed the first initial *in vivo* study and analyzed the data. F.H., Y.L., J.H.D., Y.C., K.U., and P-C.W assisted with the animal surgeries. S.H., P.N.D., J.D.B., and E.A. designed the second initial *in vivo* study and analyzed the data. D.S.A., J.H.D., D.V., H.P., J-Y.S., A.M., and A.D.D. assisted with the animal surgeries. S.H., A.M.M, J.D.B., and E.A. designed the main study experiments, analyzed the data, and wrote the manuscript. All authors collected data, and commented on and approved the final manuscript. J.D.B. and E.A. conceived and supervised the research.

## Data and materials availability

hMSCs and gelatin microparticles (E.A.), mineral-coated hydroxyapatite microparticles (W.L.M.), and custom fixation plates (J.D.B.) can be obtained through an MTA.

